# Short exon prediction based on multiscale products of a genomic-inspired multiscale bilateral filtering

**DOI:** 10.1101/423053

**Authors:** Xiaolei Zhang, Weijun Pan

## Abstract

Multiscale signal processing techniques such as wavelet filtering have proved to be particularly successful in predicting exon sequences. Traditional wavelet predictor is domain filtering, and enforces exon features by weighting nucleotide values with coefficients. Such a measure performs linear filtering and is not suitable for preserving the short coding exons and the exon-intron boundaries. This paper describes a short exon prediction framework that is capable of non-linearly processing DNA sequences while achieving high prediction rates. There are two key contributions. The first is the introduction of a genomic-inspired multiscale bilateral filtering (MSBF) which exploits both weighting coefficients in the spatial domain and nucleotide similarity in the range. Similarly to wavelet transform, the MSBF is also defined as a weighted sum of nucleotides. The difference is that the MSBF takes into account the variation of nucleotides at a specific codon position. The second contribution is the exploitation of inter-scale correlation in MSBF domain to find the inter-scale dependency on the differences between the exon signal and the background noise. This favourite property is used to sharp the important structures while weakening noise. Three benchmark data sets have been used in the evaluation of considered methods. By comparison with two existing techniques, the prediction results demonstrate that: the proposed method reveals at least improvement of 50.5%, 36.7%, 12.8%, 17.8%, 17.7%, 11.5% and 12.2% on the exons length of 1-49, 50-74, 75-99, 100-124, 125-149, 150-174 and 175-199, respectively. The MSBF of its nonlinear nature is good at energy compaction, which makes it capable of locating the sharp variations around short exons. The direct scale multiplication of coefficients at several adjacent scales obviously enhanced exon features while the noise contents were suppressed. We show that the non-linear nature and correlation-based property achieved in proposed predictor is greater than that for traditional filtering, which leads to better exon prediction performance. There are some possible applications of this predictor. Its good localization and protection of sharp variations will make the predictor be suitable to perform fault diagnosis of aero-engine.

## 1 Introduction

Recent advancement in high-throughput analysis, such as next-generation sequencing, has resulted in the development of computational techniques for the rapid prediction of exons in DNA sequences. Although great progress has been made in the development of exon prediction algorithms, the challenge of determining the lengths and locations of short exons (<200 base pair (bp)) urgently needs to be solved [1-3]. The main difficulty in predicting short exons is that the intrinsic properties, such as codon biases, are harder to determine [3,4]. In another work [5], we have briefly outlined the intrinsic advantages and limitations of the existing methods for predicting short exons. In this paper, we focus on the development of a spectral analysis technique for finding short exons in eukaryotic DNA sequences, as described below.

The discrete nature of DNA information has been driving a surging interest in the application of the principles of spectral analysis to develop efficient exon-prediction techniques. Spectral analysis techniques are attractive because they are easy to implement, entail reduced computational complexity, and mostly do not require any training of the genomic data [6-8]. In the spectral analysis of DNA sequences, the three-base periodicity (TBP) exhibited by exons is a good discriminator of coding potential. The determination of TBP due to codon usage bias is built upon the phenomenon that exon regions have a prominent power spectrum peak at frequency *f* = 1/3 [6,7]. Numerous advanced exon-finding algorithms have been developed by tracking the strength of TBP along a DNA sequence [3-19]. Such methodologies have a strong mathematical basis, including Fourier transform measures [6, 9-10], digital-filter-based methods [8,11], wavelet-based techniques [3,5,12-16] and other analysis tools [7]. Wavelets have proved highly successful in the manipulation and analysis of biomedical signals [20-26]. Among exon-finding methods, wavelet-based techniques are said to be distinctive. The examination of local variations in scale of the multiscale transform data of the sequence makes the wavelet predictor more powerful. Traditionally, the base idea of wavelet predictor, such as the modified Gabor-wavelet transform [12] and the wide-range wavelet window (WRWW) [16], is to computes a weighted sum of nucleotide values over a large neighbourhood at different scales. Although wavelet-based methods yield good predictions, they do not perform well in preserving the short exons and the exon-intron boundaries due to their linear nature.

Our intuition is that nucleotides in the codon position *p* (*p* = 1, 2, 3) are close to each other not only if they occupy nearby spatial locations but also if they have some similarity at the reading frame *p*. For this purpose, we propose a genomic-inspired multiscale bilateral filtering (MSBF) that can incorporate domain and similarity by means of multiplication. Like traditional wavelet predictor, a domain filtering named B-spline wavelet transform is designed to extract TBP by weighing nucleotide values with complex coefficients. Similarly, we define range filtering, which measures similarity by counting the sum of difference for variable sequence coverage. Another object of this paper is to investigate the inter-scale correlation information in MSBF domain and its application to short exon prediction. We formulate the problem of investigating the correlated features in terms of the differences between exon and intron coefficients at two adjacent scales. We pursue this investigation which results from the HMR195 dataset by calculating the Jensen-Shannon divergence and the histogram distributions. Experimental results demonstrate that through MSBF and multiscale products, detection accuracy can be significantly improved with only a small loss in the prediction of short exons. The proposed technique, termed multiscale products in MSBF domain (MP-MSBF), is more effective than locating exons directly from the linear filtering data, leading to superior exon prediction results.

## 2 Methodology

### 2.1 Numerical representation of a DNA sequence

The representation of DNA character strings into numerical sequences is the first step in DNA spectral analysis. In this paper, the paired-numerical representation [6] is introduced to map DNA characters (i.e., A, C, G, and T) into numeric values. A particular advantage of this representation is that it exploits the structural differences between exon and intron regions to facilitate the TBP extraction, in addition to reducing complexity. Eq (1) provides an example of this representation scheme for the short DNA fragment *…CTGCAGTGGT…*:

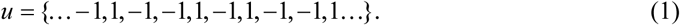

### 2.2 Multiscale bilateral filtering (MSBF)

To introduce the MSBF, we first describe in Section A the domain filtering called B-spline wavelet transform (BSWT). This wavelet function exhibits a higher degree of freedom for curve design, which can be adapted to analyse complex genome. In the next section, we first define a continuous representation of the average magnitude difference function (AMDF) inspired by [6], and a range filtering built with AMDF is designed to find certain information about nucleotide similarity in a specific codon position. Finally, the proposed MSBF is suggested for differentiating between intron noise and meaningful data.

#### A Domain filtering

In this work, wavelet functions given by B-spline windows are formulated. The B-spline window *β*_*m*_ (*t*) of order *m*, which is time-limited in [-*T*/2, *T*/2], is built as follows [27]:

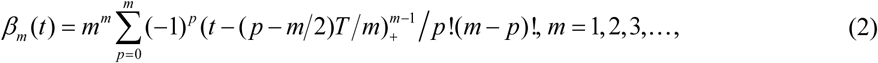

Where

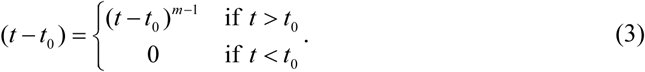

Fig. 1(A) plots *β*_*m*_ (*t*) following Eqs (2) and (3).

**Fig. 1.**
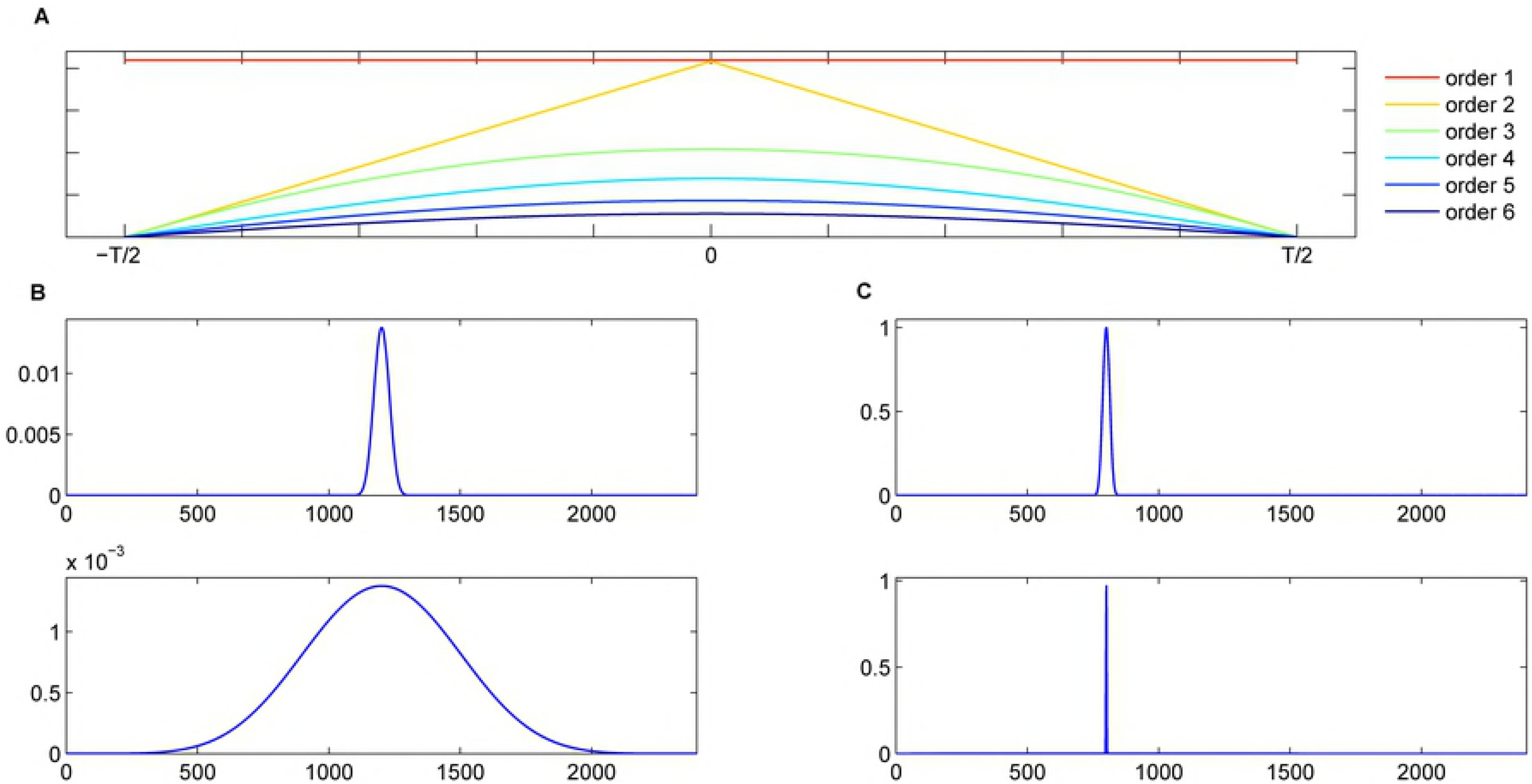
Examples of B-spline windows, and B-spline wavelet function (order 6) with two different scales in the time and frequency domains. (A) Examples of B-spline windows; (B) Magnitude response of B-spline wavelet function in the time domain; (C) Frequency response of B-spline wavelet function.

To fully analyse the DNA sequences characterized by a specific periodicity, the task here is to extract the TBP at different scales while keeping the analysis frequency constant. From Eqs (2) and (3), our proposed wavelet function of length *L* is defined as

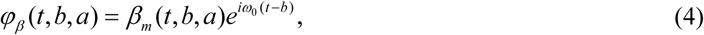

where *i*^2^ = −1, *ω*_0_ = *L/*3 denotes the basic frequency, and *β*_*m*_ (*t, b, a*) are families generated from the base functions *β*_*m*_ (*t*) by dilations and translations, i.e.,

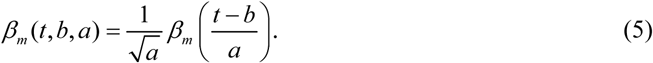

Fig. 1(B) and Fig. 1(C) illustrate our B-spline wavelet with two different scales in the time and frequency domains. The proposed BSWT of a signal *u* is given by

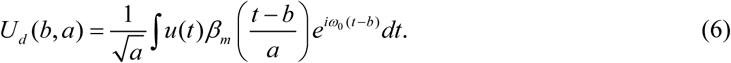

In the case of BSWT, the length of the wavelet function is 2400, and the scale parameter is set to 20 exponentially separated values between1 60 and1 6 for an input sequence. For practical purposes, the order of the B-spline function *β*_*m*_ (*t*) is truncated to 6.

#### B Range filtering

Before continuing to the proposed MSBF, we first use the AMDF to design a range filtering for measuring nucleotide similarity in a specific codon position. A continuous representation of AMDF for a signal *u*, as a function of the grid spacing 3, is defined as

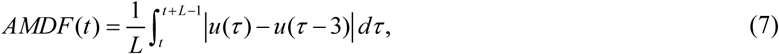

where *L* is equal to the window length of BSWT. Before applying *AMDF* to a DNA sequence, the authors in [6] suggest passing it first through a second-order resonant filter centered at frequency 2*π/* 3 [11].

For efficient implementation, a multiscale and sliding window will move along the filtered sequence to compute AMDF for the whole sequence. The expression given in Eq (4) is then used to calculate the window:

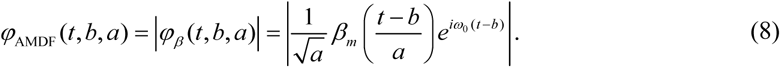

In other words, the window*φ*_AMDF_ (*t, b, a*) is the magnitude response of *φ*_*β*_(*t, b, a*) in time domain. From Eqs (7) and (8), the proposed range filtering for a signal *u* can be formulated as follows:

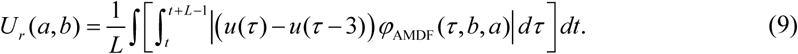

Finally, the expressions given in Eqs (6) and (9) are used to design our proposed MSBF having the non-linear property:

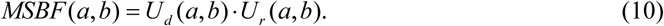

Given a DNA sequence of length *N*, the projection of the MSBF coefficients onto the position axis is defined as a function of *b* (*b* = 0,1,?, *N* −1).

### 2.3 Multiscale products

Several exon-finding techniques take advantage of traditional wavelet transform to filter short exons with small scales and long exons with large scales. This approach implies that they do not exploit the dependencies between adjacent scales. To explore the MSBF inter-scale correlations we multiply the adjacent MSBF subbands to distinguish intron noise from meaningful data while preserving the sharp variations of short exons. The core idea behind the multiscale products method is based on our research (see **Section 3.4**): namely, for DNA sequences represented by MSBF, the multiscale transform coefficients related to intron noise are less correlated across scales than are the coefficients associated with exon signals.

Let *U* (*a*_*j*_, *b*) be the MSBF of a signal *u* at the scale *a* _*j*_ (*j* = 1, 2,… *J*) and the position *b*. The multiscale products (or inter-scale correlation) *MP*(*a*_*j*_, *b*) of the MSBF contents at two adjacent scales is defined as

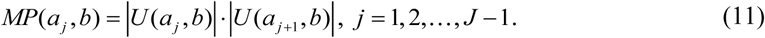

With the observation of experimental results, we can imagine that multiplying the MSBF at adjacent scales would amplify exon structures and dilute noise (see Section 3).

### 2.4 Procedure of MP-MSBF predictor

Our MP-MSBF method for exon prediction is described briefly in Table 1. The input DNA sequence of length *N* is referred to as *u*.

**Table 1.**
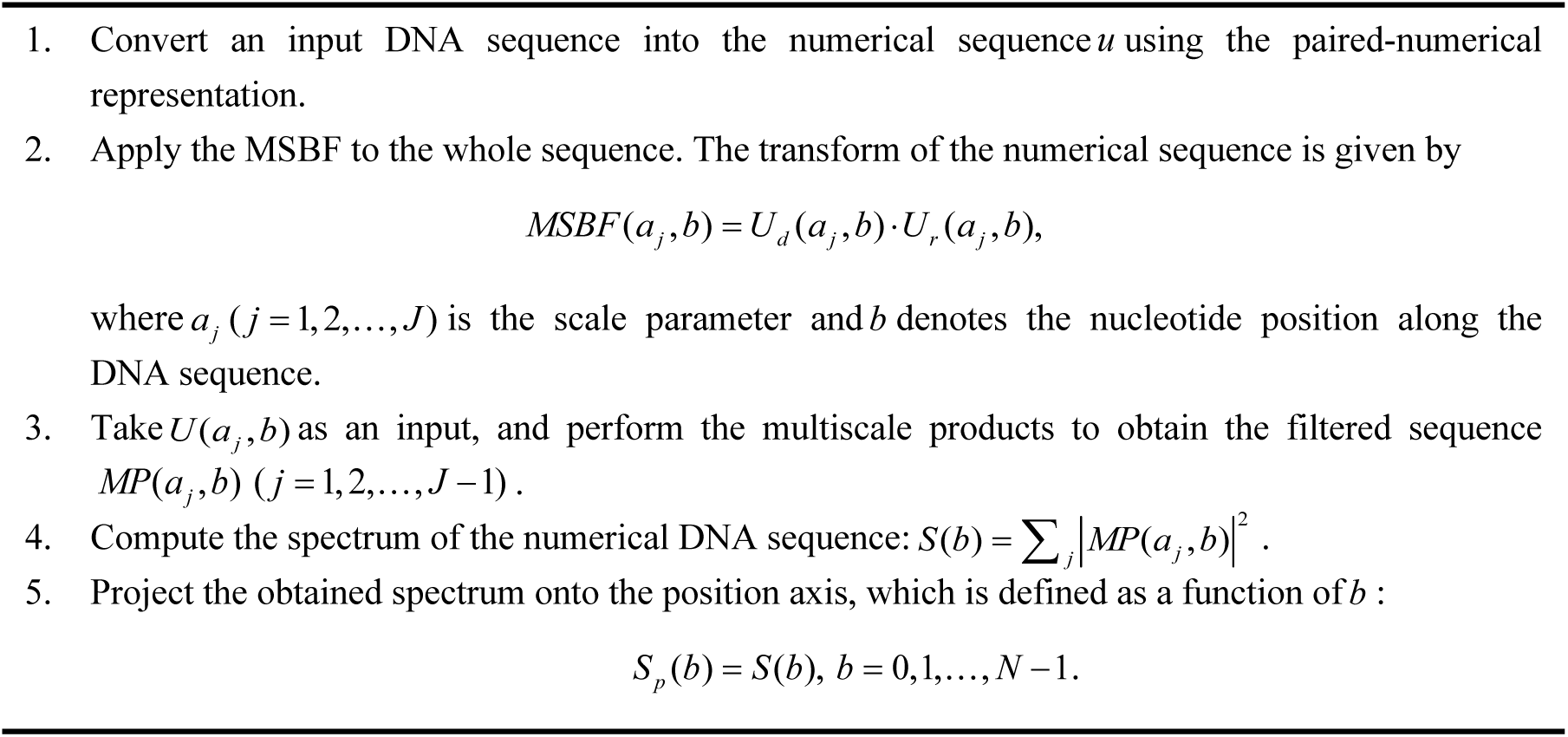
Exon predictor algorithm using the MP-MSBF technique.

## 3 Results and discussion

### 3.1 Data resources

To evaluate and compare the performance of the proposed MP-MSBF with that of other methods, the two benchmark data sets BG570 [28] and HMR195 [29] have been considered. Furthermore, we conduct an additional classification experiment using 29 sequences of the ENm001 and ENm004 data sets (part of EGASP) [30] (see **S1 File** for detailed information on these sequences). Table 2 summarizes the features of the considered data sets.

**Table 2.** Statistics of the test data sets.

| Dataset | Species | Genes | Length | Exons | Average length of exons | Proportion of exons | Exons (<200) density in total exons |
| --- | --- | --- | --- | --- | --- | --- | --- |
| BG570 | Vertebrate | 570 | 2,892,149 | 2,649 | 168 | 15.37% | 78.86% |
| HMR195 | Mammalian | 195 | 1,383,720 | 948 | 208 | 14% | 77.43% |
| EGASP | Human | 29 | 2,425,886 | 893 | 157 | 5.8% | 77.83% |

### 3.2 General setting

In this section, we first conduct an experiment to establish a comprehensive analysis of the inter-scale correlation of the differences between exon and intron coefficients. Next, we present experiments in exon prediction using the proposed method. For comparison, MP-MSBF presents comparable performance to that of two popular existing methods: fast Fourier transform plus empirical mode decomposition (FFTEMD) [9] and WRWW [16]. To evaluate the general performances of these measures, the TBP data for each DNA sequence considered have been normalized to [0,1].

### 3.3 Evaluation metrics

To investigate the inter-scale correlation of the differences between exon and intron sequences, the distance criterion of Jensen-Shannon (JS) divergence [31] is adopted. In probability theory and statistics, the JS divergence is a method of measuring the similarity between two probability distributions. The JS divergence is a convenient divergence measure for our purpose because it is symmetric and bounded between 0 and 1. The distance between two probability vectors ***P*** and ***Q*** in terms of the JS divergence is defined as

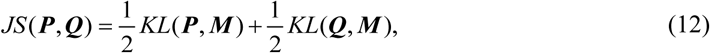

Where ***M*** = (***P*** + ***Q***) 2 and *KL* is the Kullback-Leibler divergence,

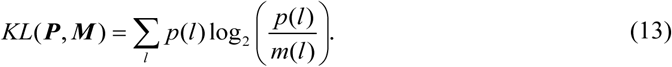

With a set of results obtained by running a predictor on a test data set, the true positive (TP), true negative (TN), false negative (FN) and false positive (FP) counts can be determined. Using these counts, the performances of various methods in handling exons of different lengths are measured in terms of the approximate correlation (AC) [28]

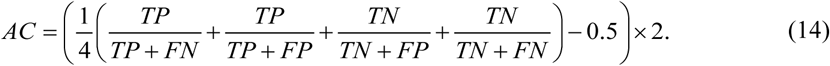

To evaluate the general performance of the method under consideration, the receiver operating characteristic (ROC) curve [32] is used to explore the effects on *sensitivity* and *specificity*. The sensitivity and specificity are given by

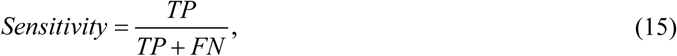

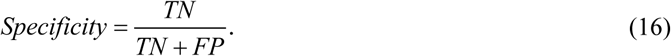

The area under an ROC curve (AUC) can be used as an indicator of prediction performance.

### 3.4 Inter-scale correlation analysis

The coefficients of the input DNA sequences obtained from the multiresolution decomposition include exon-structure information together with intron noise. The general purpose of inter-scale correlation analysis is to investigate the dependency information on the differences between exon and intron coefficients. We apply the schemes proposed in this paper to analyse the correlation for a large number of exon and intron regions.

Fig. 2 shows the prediction plots of the sequence HUMDZA2G of *Homo sapiens* (GenBank accession number D14034) using MSBF and its inter-scale correlation (or multiscale products) at different scales. The sequence HUMDZA2G contains four exons at positions 5322-5388, 9329-9589, 12907-13182 and 14052-14335. The peaks corresponding to the exon regions of the original data appear much stronger in Fig. 2(B) than in Fig. 2(A). The results demonstrate that inter-scale correlation can suppress intron noise while retaining more exon details. Fig. 3 compares the prediction results of the sequence HUMDZA2G using the tested methods. Our MP-MSBF algorithm identified the localized peaks better and located the short coding sequence (exon 1) more accurately.

**Fig. 2.**
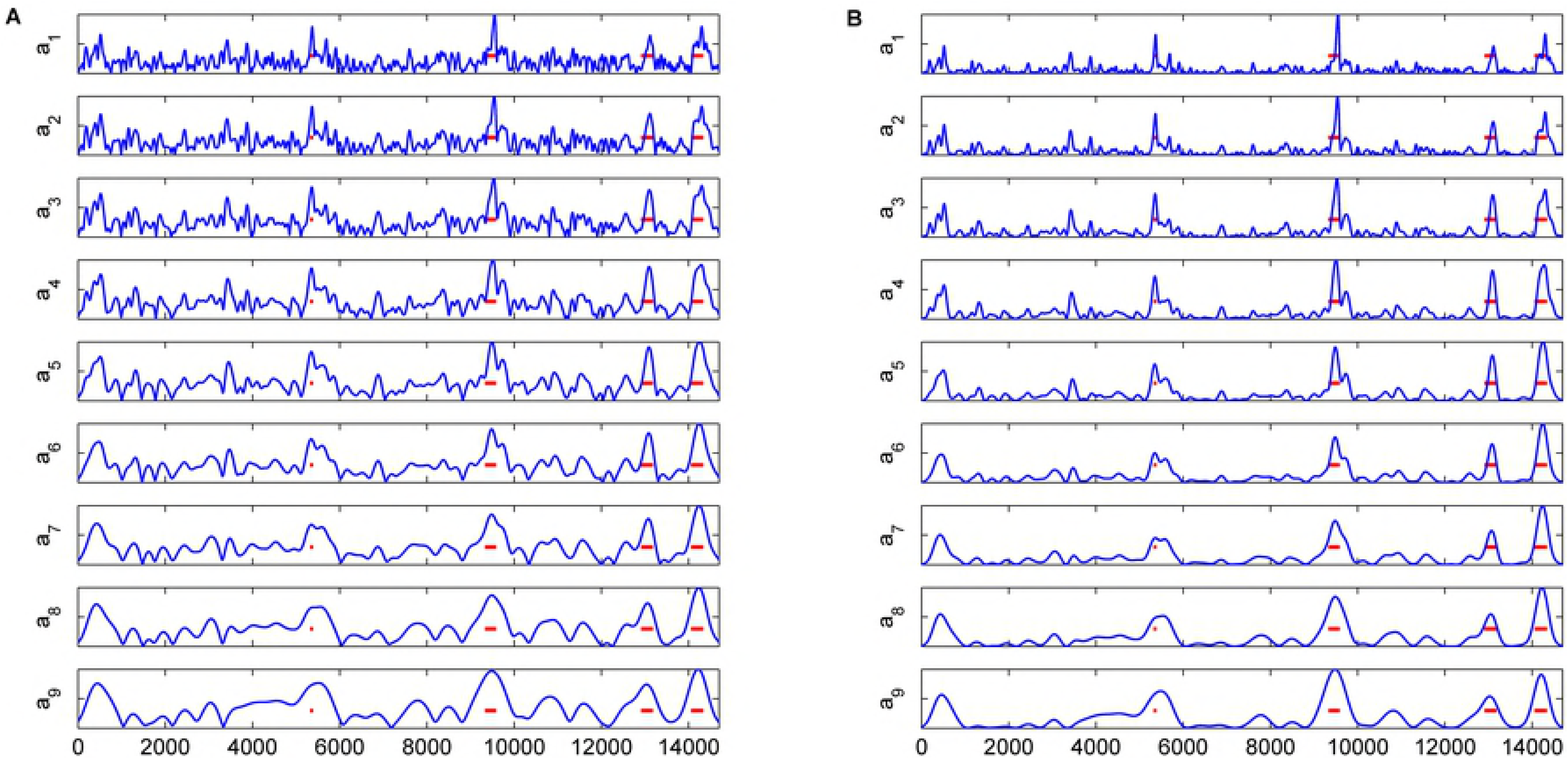
Prediction plots for sequence HUMDZA2G at different scales. The abscissa axes of all the plots represent the relative base positions, and the actual locations of the exons are marked with red horizontal lines. Part (A) shows the MSBF result; and (B) shows the result of inter-scale correlation.

**Fig. 3.**
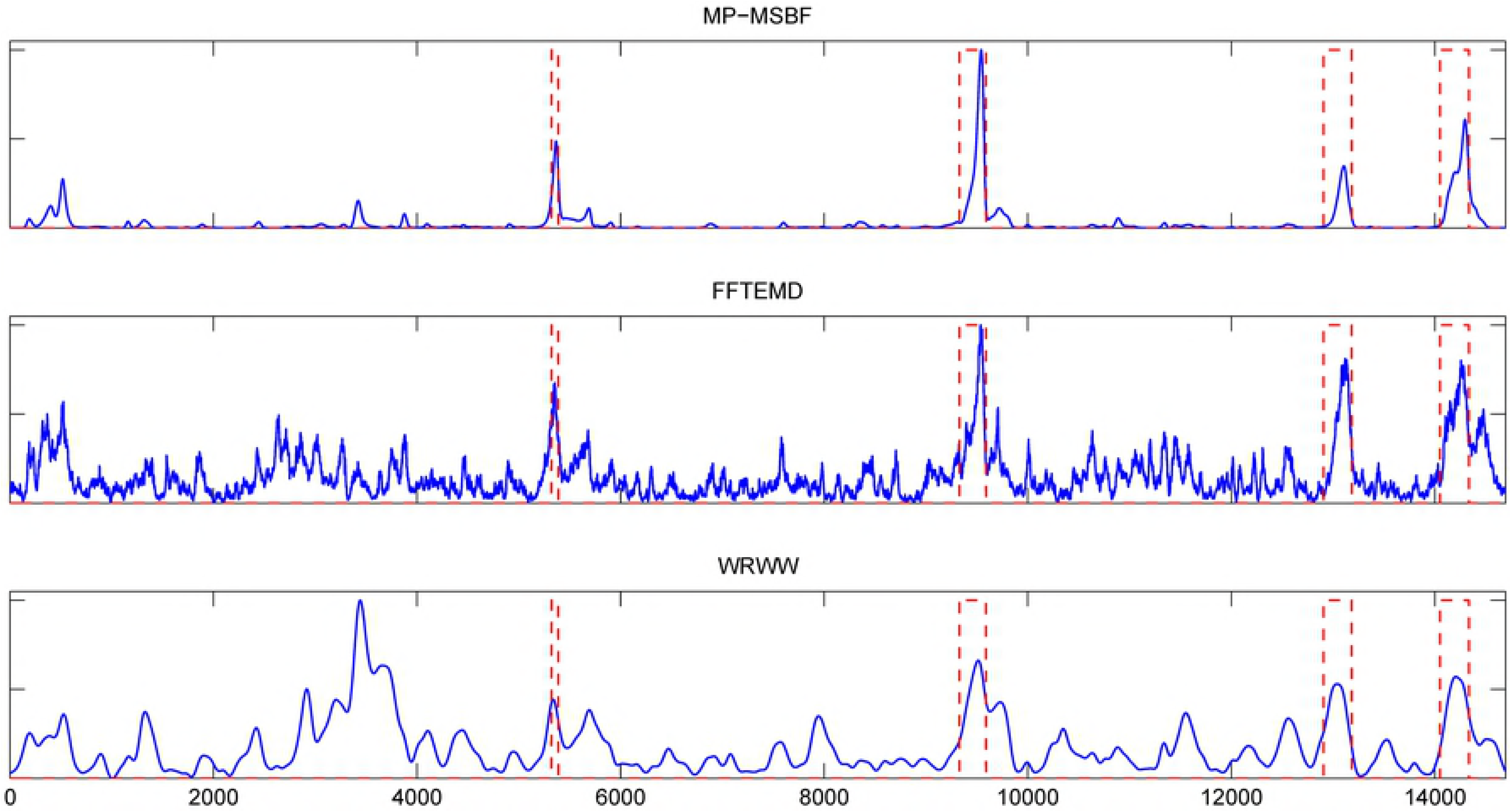
Prediction results for the sequence HUMDZA2G using the considered methods. The abscissa axes of all the plots represent the relative base positions, and the actual locations of the exons are marked with rectangles in red dashed lines.

Herein, the JS divergences are employed to investigate whether the coefficients related to introns are less correlated across scales than the coefficients associated with exons. This distance criterion has been applied in genome comparison [33], bioinformatics [34] and protein surface comparison [35]. Table 3 summarizes the JS divergences of the MSBF coefficients between two adjacent scales, *a*_*j, j* +1_ (*j* = 1, 2,…, 9), for the exon and intron nucleotides of the HMR195 data set. The results of Table 3 reveal that the JS divergences of exons are smaller than those of introns at consecutive scales, while there is a difference of one order of magnitude between the exon and intron regions at the last eight consecutive scales. This property can assist in discriminating short exon features from introns in the multiscale transform domain.

**Table 3.** JS divergence of MSBF coefficients between two adjacent scales.

| Regions | JS divergence of MSBF coefficients between consecutive scales |  |  |  |  |  |  |  |  |
| --- | --- | --- | --- | --- | --- | --- | --- | --- | --- |
| | $a_{1,2}$ | $a_{2,3}$ | $a_{3,4}$ | $a_{4,5}$ | $a_{5,6}$ | $a_{6,7}$ | $a_{7,8}$ | $a_{8,9}$ | $a_{9,10}$ |
| Exon | 0.0030 | 0.0028 | 0.0032 | 0.0057 | 0.0083 | 0.0089 | 0.0041 | 0.0035 | 0.0033 |
| Intron | 0.0085 | 0.0130 | 0.0189 | 0.0220 | 0.0206 | 0.0138 | 0.0102 | 0.0133 | 0.0178 |

To further justify our assumption, histograms with fitted distributions are calculated for the exon and intron nucleotides of HMR195 at different scales. Fig. 4 and Fig. 5 give the distributions of exon and intron nucleotides using the MSBF and inter-scale correlation, respectively. This result indicates that the most relevant exon information represented by the correlation at each scale is captured by large-valued coefficients, whereas the intron information is captured by a large number of small-valued coefficients. Fig. 6 clearly illustrates that the distance between the exon and intron curves obtained from inter-scale correlation is greater than that obtained from MSBF. In other words, the MSBF coefficients of the exon sequences have a strong correlation on various decomposition scales, whereas the MSBF coefficients of noise are weakly correlated. These plots justify our assumption.

**Fig. 4.**
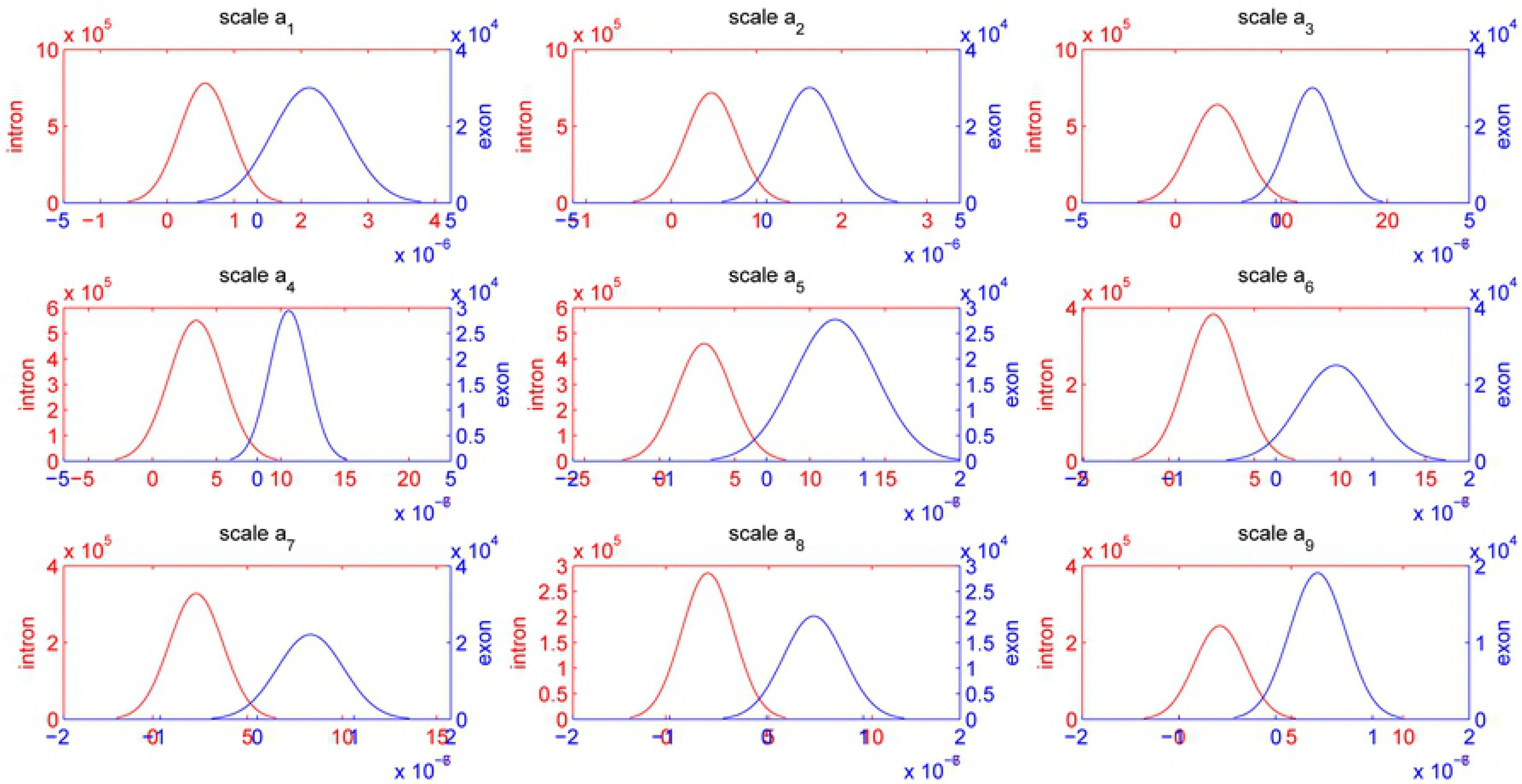
Histogram distributions at different scales for MSBF applied to HMR195. For all the plots, blue lines represent exons, red lines indicate introns, the abscissa axes represent the magnitude values, and the ordinate axes represent the number of MSBF coefficients.

**Fig. 5.**
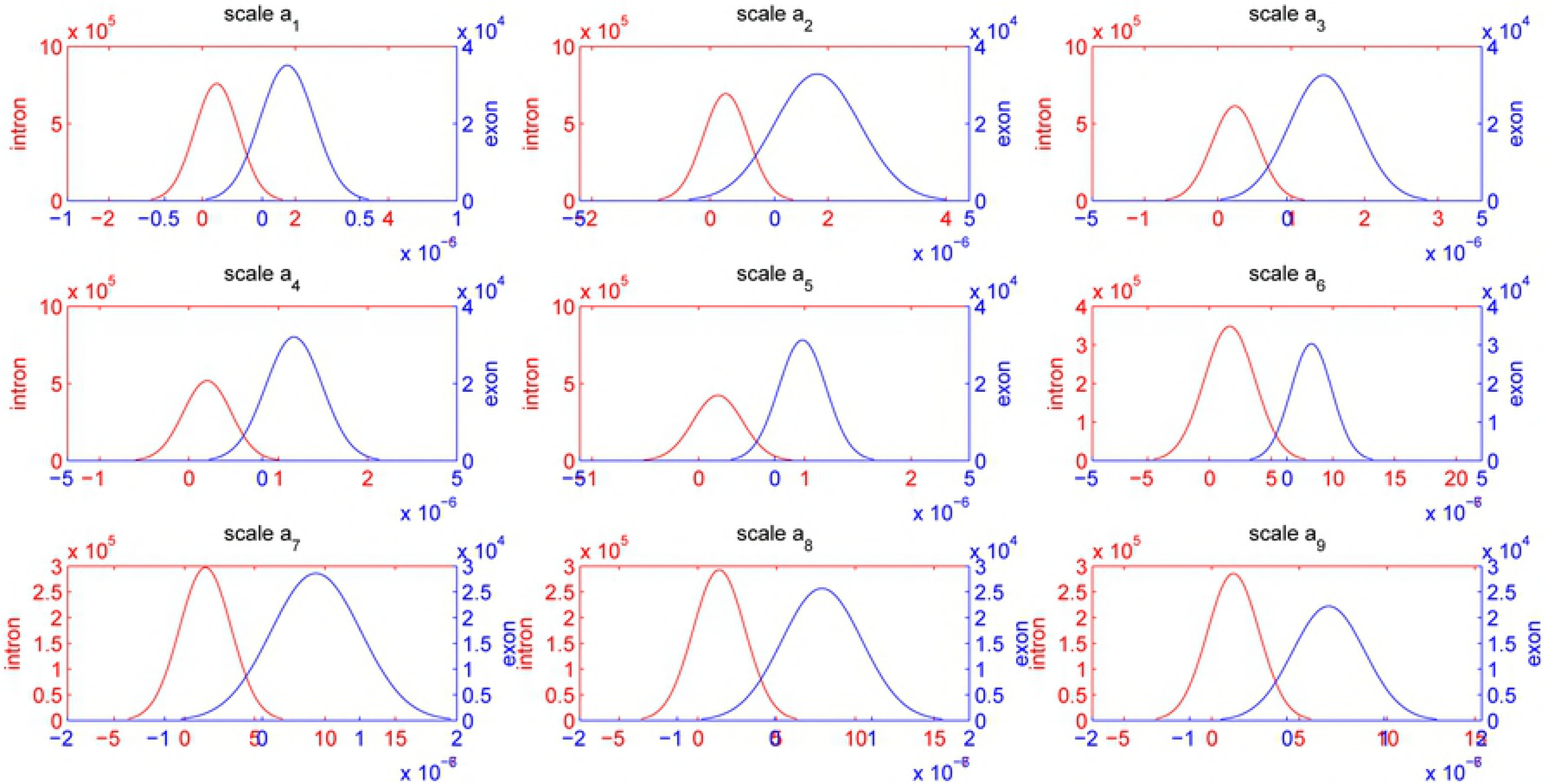
Histogram distributions at different scales for inter-scale correlation applied to HMR195. ‡ See Fig. 4 for legend.

**Fig. 6.**
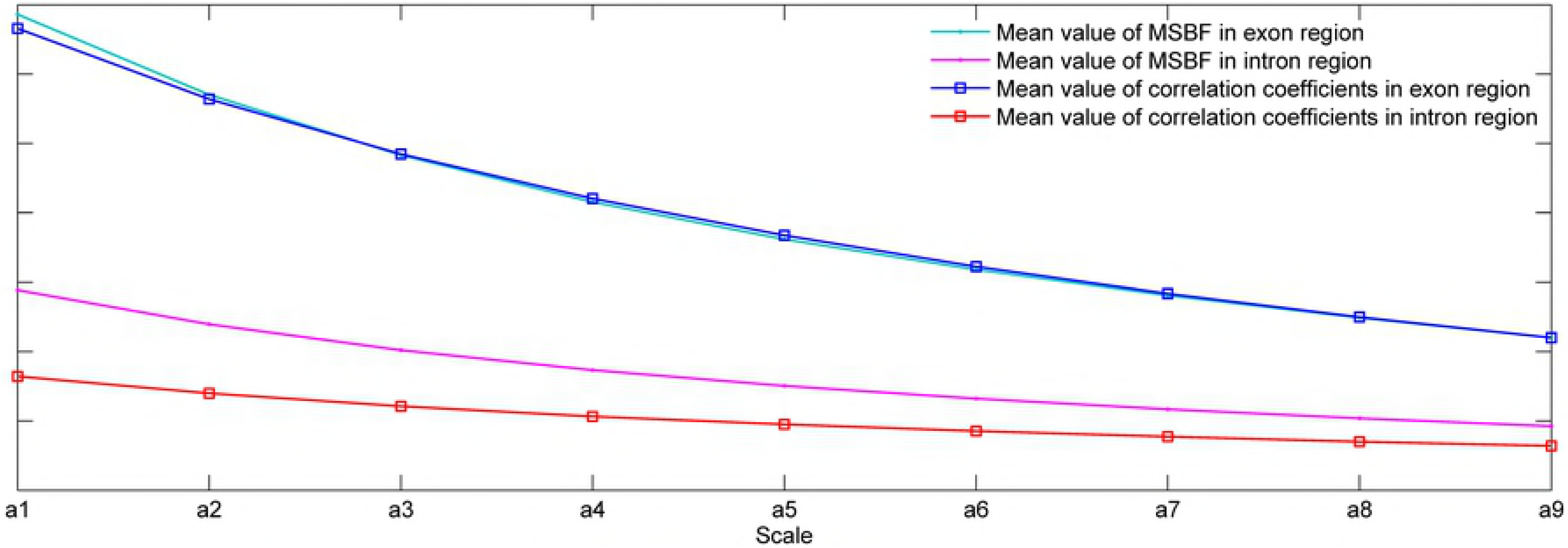
Mean values of histogram distributions at different scales for MSBF and inter-scale correlation applied to HMR195.

† See Fig. 4 for legend.

### 3.5 Performance evaluation on benchmark data sets

Exons have significant functional constraints, and their length plays an important role in splice site selection. Rogic’s evaluation work [29] stated, “*These constraints have shaped the exon length distribution quite differently from geometric distribution. The length distribution depends on the exon type*.” In our analysis, we grouped exons into eight ranges of exon lengths, namely, (0,50), [50,75), [75,100), [100,125), [125,150), [150,175), [175,200), [200,300) and [300,300+). The exons within the range (0,200) are considered small [2]. The best accuracies achieved by the tested methods are calculated in terms of the AC values for each group of exons. Fig. 7(A) and Fig. 7(B) depict the experimental results obtained from various methods using the BG570 and HMR195 data sets, respectively. The results show that the MP-MSBF outperforms the other tested methods in these nine ranges of exon lengths. Similar results obtained with the sequences in the ENm001 and ENm004 data sets are shown in Fig. 7(C). The MP-MSBF exhibits good accuracies in these nine ranges and presents results close to those of WRWW in the ranges [50,75), [100,125) and [200,300), while it slightly exceeds the performance of other methods in the other ranges. Fig. 7 shows that all methods tend to have low accuracy in predicting exons shorter than 50. A possible explanation for this phenomenon is that this type of exon is too short to be efficiently spliced in vivo without special splicing activation sequences [36]. For almost all the methods, the accuracies slowly rise with the length of annotated exons between 75 and 200 nucleotides.

**Fig. 7.**
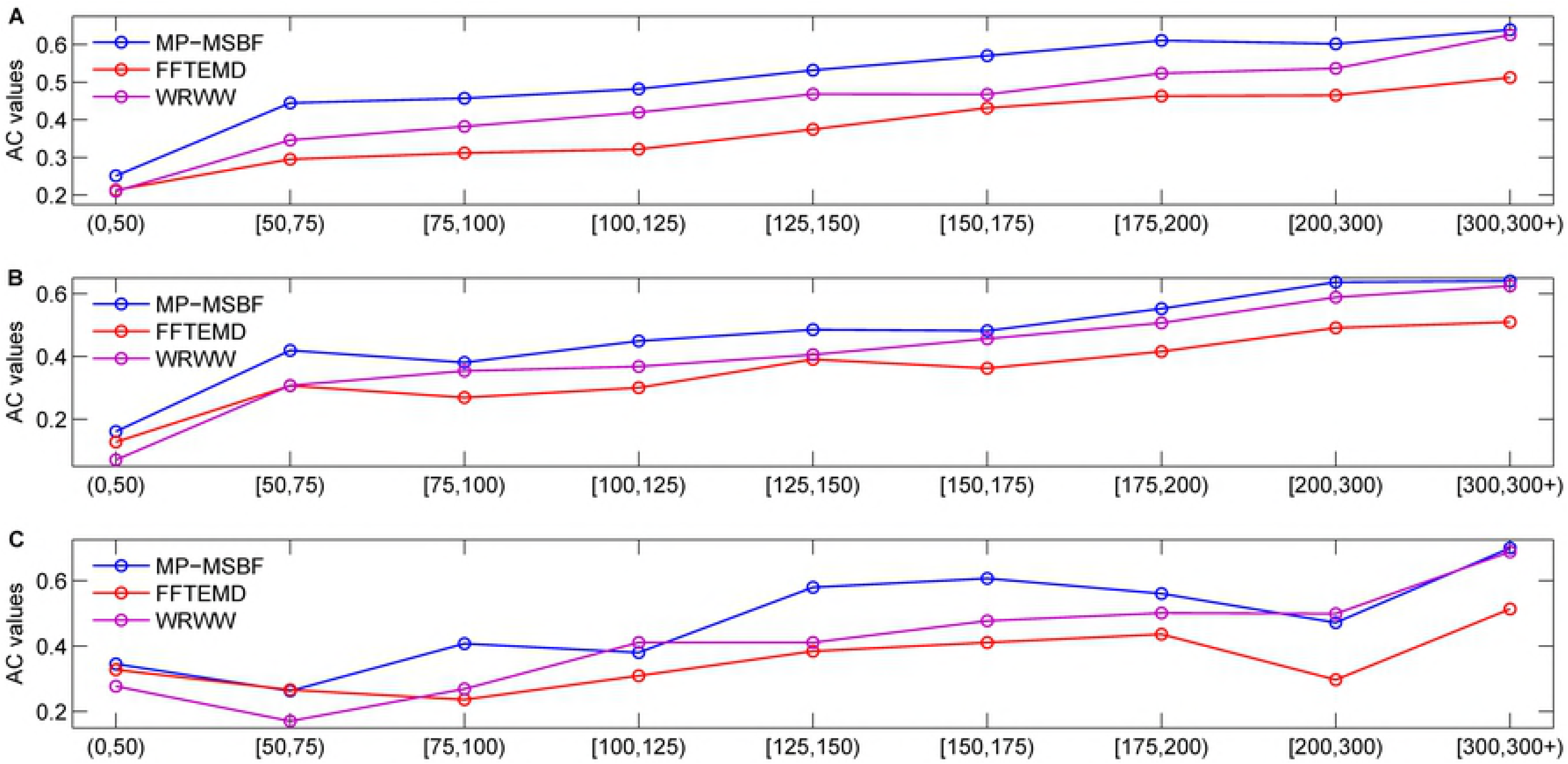
Plots of AC for considered data sets with various methods applied to exons in nine length ranges. For all the plots, the ordinate axes denote the ranges of exon lengths.

Table 4 summaries the performances of various methods for short exons (<200) using the BG570, HMR195, ENm001 and ENm004 data sets. By comparison with the FFTEMD and WRWW methods, the prediction results show that: our MP-MSBF exhibits at least improvement of 50.5%, 36.7%, 12.8%, 17.8%, 17.7%, 11.5% and 12.2% on the exons of the ranges (0,50), [50,75), [75,100), [100,125), [125,150), [150,175) and [175,200), respectively.

**Table 4.** Best performances obtained from considered methods for short exons using the BG570, HMR195, ENm001 and ENm004 data sets.

| Methods | Approximate coefficient (AC) |  |  |  |  |  |  |
| --- | --- | --- | --- | --- | --- | --- | --- |
|  | (0,50) | [50,75) | [75,100) | [100,125) | [125,150) | [150,175) | [175,200) |
| MP-MSBF | 0.2687 | 0.4657 | 0.4593 | 0.5050 | 0.5656 | 0.5630 | 0.6228 |
| FFTEMd | 0.1785 | 0.3330 | 0.3335 | 0.3520 | 0.4389 | 0.4238 | 0.4754 |
| WRWW | 0.1598 | 0.3406 | 0.4073 | 0.4289 | 0.4807 | 0.5051 | 0.5551 |

An additional classification experiment on all sequences of considered data sets is designed to assess the general performance of our proposed technique and other methods. Fig. 8 presents the ROC curves obtained from the different methods tested in this experiment. The MP-MSBF method has higher prediction accuracy than its counterparts. Our MP-MSBF method consistently exhibits higher prediction accuracy than its counterparts for exons that are either relatively short or long in length.

**Fig. 8.**
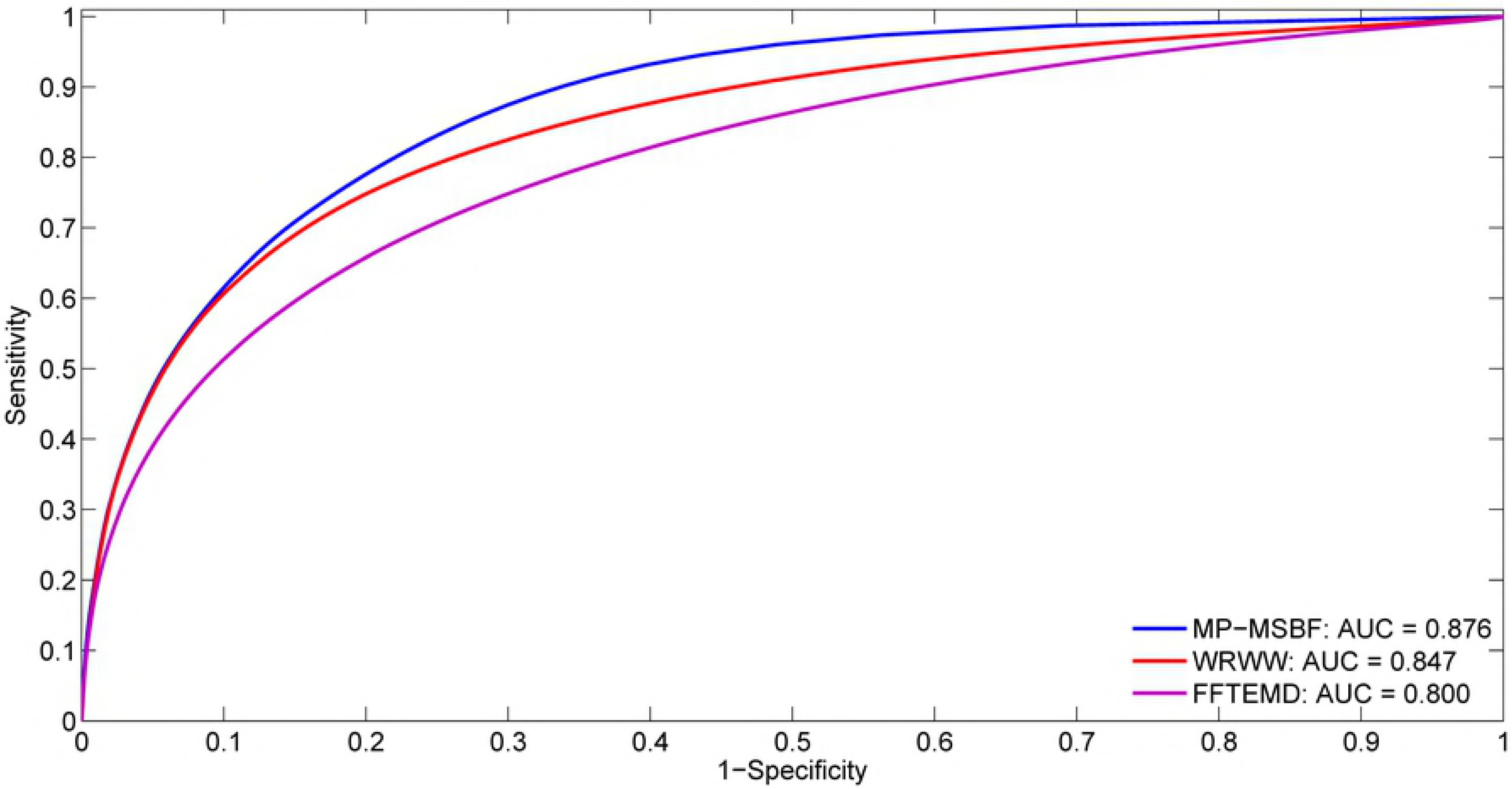
ROC plots of tested methods using the BG570, HMR195, ENm001 and ENm004 data sets.

### 3.6 Summary

In this work, we have introduced a new, robust and efficient method to predict exons in eukaryotes. Unlike some prediction techniques that detect exons directly by linear filtering, the proposed scheme incorporates a genomic-inspired multiscale bilateral filtering and its inter-scale dependencies and then applies these features to better differentiate exon structures from background noise. The first key concept of our method is its nonlinear nature, which exploits geometric distance in the spatial domain and nucleotide similarity in the range. The second is that this technique compacts the energy of the exons into coefficients with large amplitudes and spreads the energy of the introns over a large number of coefficients with small amplitudes. This phenomenon has led to improved results with respect to short exon preservation and noise suppression. The proposed MP-MSBF method requires neither prior information nor training models for exon prediction, and so it can be applied to analyse unknown and novel genomes. Although not good for exons shorter than 50, the results obtained from our method are acceptable.

## 4 Conclusion

Exons encode the biochemical processes and information involved in the pathway from DNA to proteins. In genomic sequence analysis, exon prediction based on the annotated sequences in the online databases is an important problem. For exon prediction, extracting the relevant features of short coding sequences is a major task because the subtle features of short exons are obscured by the strong presence of background noise. In practice, spectral analysis is an important tool for the discovery of interesting patterns and structures in short exon data. In this paper, we present a new exon-finding spectral analysis method that overcomes some of the shortcomings of current predicting techniques. The MP-MSBF predictor takes advantage of the nonlinear filtering and the dependency information between scales, which makes it capable of short exon prediction. We see some possible applications of this predictor. The correlation-based property and nonlinear nature of this technique allow the selection of a characteristic frequency from surrounding noise and thereby makes it possible to offer good localization and protection of sharp variations for locating hot spots in proteins and performing fault diagnosis of aero-engine.

## Supporting information

**S1 File. Sequences of the ENm001 and ENm004 data sets used for the analyses presented in this paper.** Detailed information on these sequences. (ZIP)

## Acknowledgments

This work was sponsored by the National Natural Science Foundation of China (Grant No. U1733203).

## Author contributions

**Conceptualization:** XLZ.

**Data curation:** XLZ.

**Formal analysis:** XLZ.

**Funding acquisition:** WJP.

**Investigation:** XLZ WJP.

**Methodology:** XLZ.

**Project administration:** XLZ WJP.

**Resources:** XLZ WJP.

**Software:** XLZ.

**Supervision:** XLZ WJP.

**Validation:** XLZ WJP.

**Visualization:** XLZ.

**Writing - original draft:** XLZ.

**Writing - review & editing:** WJP.

